# Genotoxic formaldehyde and lipid aldehydes are sources of DNA damage in keratinocytes

**DOI:** 10.1101/2025.11.13.688345

**Authors:** Nicolas J. Blobel, Yibing Yao, Marian C. Okondo, Agata Smogorzewska

## Abstract

The Fanconi anemia (FA) pathway is vital for the repair of DNA interstrand crosslinks (ICLs), which are caused by a variety of endogenous and exogenous genotoxins including reactive aldehydes. Patients with pathogenic variants in the FA pathway are predisposed to early-onset, aggressive malignancies, especially leukemia and head and neck, esophageal, and anogenital squamous cell carcinomas (SCCs). Prior studies have linked endogenous formaldehyde and acetaldehyde in hematopoietic stem cells with bone marrow failure and leukemia in FA-deficient mice. However, the genotoxic aldehydes specific to mucosal keratinocytes, precursors of FA-associated SCCs, remain to be identified.

Here, to identify alcohol dehydrogenases (ADHs) and aldehyde dehydrogenases (ALDHs) necessary for protection of keratinocyte genomes from endogenous metabolites, we used a sensitized background of FA-pathway deficiency. We systematically inactivated all highly expressed ADH and ALDHs genes in *FANCA*-deficient keratinocyte cell lines and identified genes required for their survival. We report that loss of ADH5 or ALDH3-family enzymes in FA pathway-deficient cells has a synthetic lethal effect and induces DNA damage markers, nominating these genes as important defense mechanisms in the detoxification of endogenous aldehydes and prevention of carcinogenesis in keratinocytes. Loss of ADH5 increases formaldehyde levels, while simultaneous loss of four functionally redundant ALDH3 isozymes is predicted to cause accumulation of lipid aldehydes. FA-deficient keratinocytes are more sensitive to treatment with formaldehyde and the lipid aldehyde 4-hydroxynonenal (4HNE) compared to FA-competent cells. In addition, the thiol-rich antioxidant N-acetyl-L-cysteine (NAC) partially rescued the growth of *FANCA^−/−^ ADH5^−/−^* cells and lowered the level of endogenous DNA damage. This work identifies important defense mechanisms in keratinocytes and suggests cancer preventive strategies in patients with Fanconi anemia.

**Significance:** This work shows that ADH5 and ALDH3-family enzymes function as tier 1 defense mechanisms against endogenous DNA-damaging metabolites in keratinocytes. Enhanced detoxification of endogenous aldehydes might prevent DNA damage and mutagenesis, which drive Fanconi anemia-associated head and neck and other mucosal squamous cell carcinomas.

## Introduction

The Fanconi anemia (FA) DNA repair pathway repairs DNA interstrand crosslinks (ICLs), lesions that covalently link the Watson and Crick DNA strands (*1*). Individuals with mutations in any of the 23 *FANC* protein-coding genes necessary for ICL repair develop Fanconi anemia, although mutations in *FANCA* are the most common (*2–8*).The spectrum of phenotypes in Fanconi anemia patients is broad and includes developmental abnormalities, bone marrow failure, infertility, and profound cancer predisposition, primarily to acute myeloid leukemia and mucosal squamous cell carcinomas (SCCs) including head and neck, esophageal and anogenital (*2, 5, 6, 8, 9*). FA patients are hundreds to thousands-fold more likely to develop early-onset mucosal SCCs and have worse clinical outcomes compared to the general population (*2, 9, 10*). With advancements in hematopoietic stem cell transplantation but limited effective cancer treatment options for FA patients, mucosal SCC has become the leading cause of death in FA patients (*11, 12*).

Genomic analysis of mucosal SCCs revealed the presence of high numbers of copy number alterations including amplifications of multiple oncogenes and deletions of tumor suppressor genes, all occurring in the setting of somatic p53 deficiency (*10*). The outcome is high genomic heterogeneity arising from the inappropriate repair of ICLs. However, the sources of the ICLs in mucosal keratinocytes are unknown. We posit that knowledge of these sources may, at least partially, explain the predisposition of patients with FA to mucosal SCCs and offer clues of preventive measures. Since we believe that the FA pathway is also important in preventing sporadic mucosal SCCs, knowledge of contributing endogenous keratinocyte-specific aldehydes may also help prevent non FA-associated cancers.

Previous work has implicated endogenous aldehydes, genotoxic metabolites that form DNA interstrand crosslinks, in the pathogenesis of bone marrow failure and leukemia in Fanconi anemia mouse models and patients (*13–19*). In particular, endogenous formaldehyde has emerged as the prime DNA-damaging metabolite that, if not detoxified, is highly deleterious to hematopoietic stem cells. Without enzymes that detoxify formaldehyde, specifically ALDH2 and ADH5, hematopoietic cells die or transform into leukemia (*13–19*). ALDH9A1 has also been shown to be important for survival of Fanconi anemia-deficient cells, implicating aminoaldehydes and their metabolites in cellular functions of hematopoietic cells and epithelial cells of the ovary, suggesting tissue specificity of DNA damaging metabolites (*20*).

Here, we aimed to identify genotoxic metabolites that might be specific to keratinocytes, the precursor cells of SCCs. Using human and mouse isogenic keratinocytes, deficient and proficient for FA pathway function and a fluorescence-based competition assay, we assessed the importance of all ALDH or ADH genes expressed in keratinocytes to protect this cell type. We found that absence of ADH5 or enzymes of the ALDH3-family decreased the survival of the FA-deficient cells through induction of DNA damage. FA-deficient keratinocytes were more sensitive to formaldehyde and lipid aldehydes than FA-competent cells. Our results expand on the proposed model of protection against endogenous metabolites in cells identifying critical drivers of DNA damage in keratinocytes as potential targets for preventing the tumorigenic transformation.

## Results

### Generation of Fanconi anemia keratinocyte cell lines

We set out to understand the sources of aldehydes that induce genome instability contributing to malignant transformation of keratinocytes. We modeled the tissue origin of most FA-associated SCCs by choosing three non-transformed human cell lines derived from primary keratinocytes of the oral cavity (**Table 1**). As we have previously discovered important differences between human and mouse aldehyde clearing enzyme expression, we have also used keratinocytes derived from a neonatal FA-deficient mouse, which we have studied in the context of carcinogenesis (*10, 20*).

For each human line, we generated FA-deficient cells lacking *FANCA*, the most mutated gene in patients with FA, using sgRNA against human *FANCA* gene (*FANCA^−/−^*) or a random intragenic region (control, *FANCA^+/+^). Fanca^−/−^* mouse keratinocytes were derived from a *Fanca^−/−^* mouse. To show functional deficiency of the FA pathway, we assessed FANCD2 monoubiquitination after treatment with the potent ICL-inducing agent mitomycin C (MMC). MMC treatment induced robust monoubiquitination of FANCD2 as assessed by immunoblotting, whereas FA-deficient cells lacked monoubiquitinated FANCD2 at baseline and failed to induce ubiquitination upon MMC treatment (**Figure 1A**). The dose-dependent sensitivity of FA-deficient cells to MMC confirmed the loss of the FA pathway, which was improved upon complementation with human *FANCA,* mouse *Fanca* cDNA and by other experiments (**Figure S1A and B** and see below).

**Figure 1.**
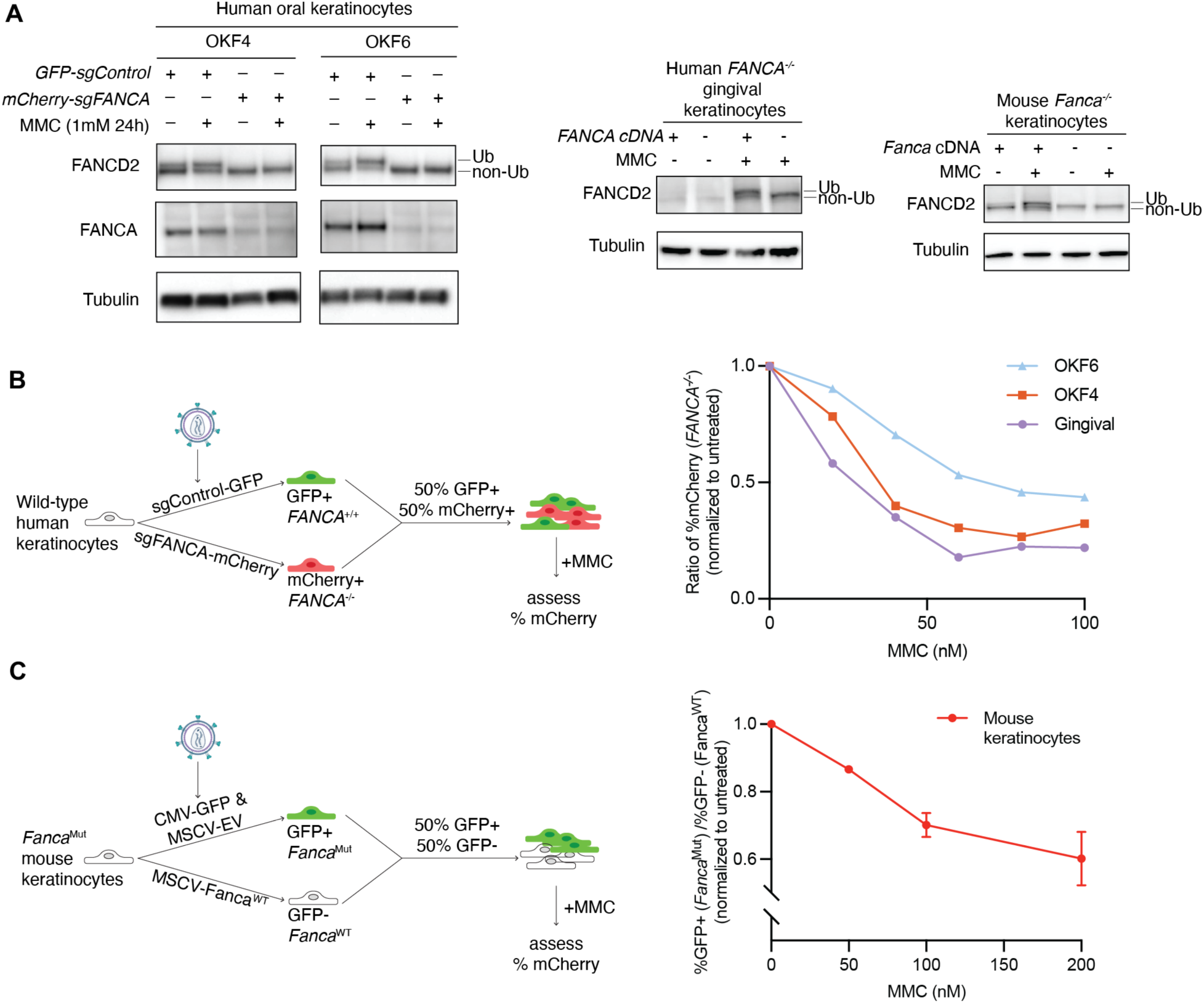
Generation and characterization of Fanconi anemia pathway-deficient keratinocyte cell lines from human and mouse. (A) Immunoblotting of FANCA and FANCD2 in FA-deficient and FA-competent human oral keratinocytes (OKF4, OKF6; left), gingival (EH; middle) and mouse keratinocytes (right), 24 hours after treatment with 1µM MMC. The monoubiquitinated form of FANCD2 is indicated as “Ub”. α-Tubulin is used as loading control. (B) MMC sensitivity of FA-deficient human keratinocytes assessed by a fluorescence-based competition assay. *Left*: Schematic of the competition assay setup. *Right*: Quantification of competition assay and the relative abundance of FA-deficient cells after treatment with increasing concentrations of MMC, normalized to the untreated control. (C) MMC sensitivity of FA-deficient mouse keratinocytes assessed by competition assay

### Validation of a competition assay between FA-deficient and FA-competent keratinocytes

One of the expectations from our previous work on identification of sources of endogenous DNA damage was that FA-deficient cells, when challenged with increased levels of such damage, exhibit progressive yet modest cell growth defects that typically require several weeks of culture to be detected (*20*). However, we have shown that competition assays between two cell populations have been a very sensitive method to discern differences in growth in such situations (*20, 21*). Thus, we first set up a competition assay between human *FANCA*^−/−^ cells labeled with mCherry and *FANCA*^+/+^ cells labeled with GFP (**Figure 1B**) or mouse GFP-labeled *Fanca^−/−^* cells and non-labeled *Fanca^+/+^* cells (**Figure 1C**). We then validated the system using MMC and observed that the percent of FA-deficient cells decreased with increasing doses of MMC treatment in all four cell lines, as expected (**Figure 1B and C**). Various iterations of this assay were used in the rest of the work to compare growth of different cells.

### *ADH5* knockout causes growth defects in FA-deficient human and mouse keratinocytes

In mammalian cells, aldehydes are primarily detoxified by 19 aldehyde dehydrogenases (ALDHs) and 8 alcohol dehydrogenases (ADHs) (*22*). To start identifying the ALDH and ADH enzymes involved in detoxification of metabolites that may be deleterious to keratinocytes deficient in the FA pathway, we first profiled the expression of these genes in the human OKF4 and OKF6 cell lines using RNA-Seq (**Figure S1C and Figure 2A**). *ALDH7A1*, *ADH5*, *ALDH9A1* are the most highly expressed aldehyde-metabolizing enzymes in these two cell lines, all with >50 Transcripts Per Million (TPM). 12 of them had detectable expression, of more than 2 TPMs.

**Figure 2.**
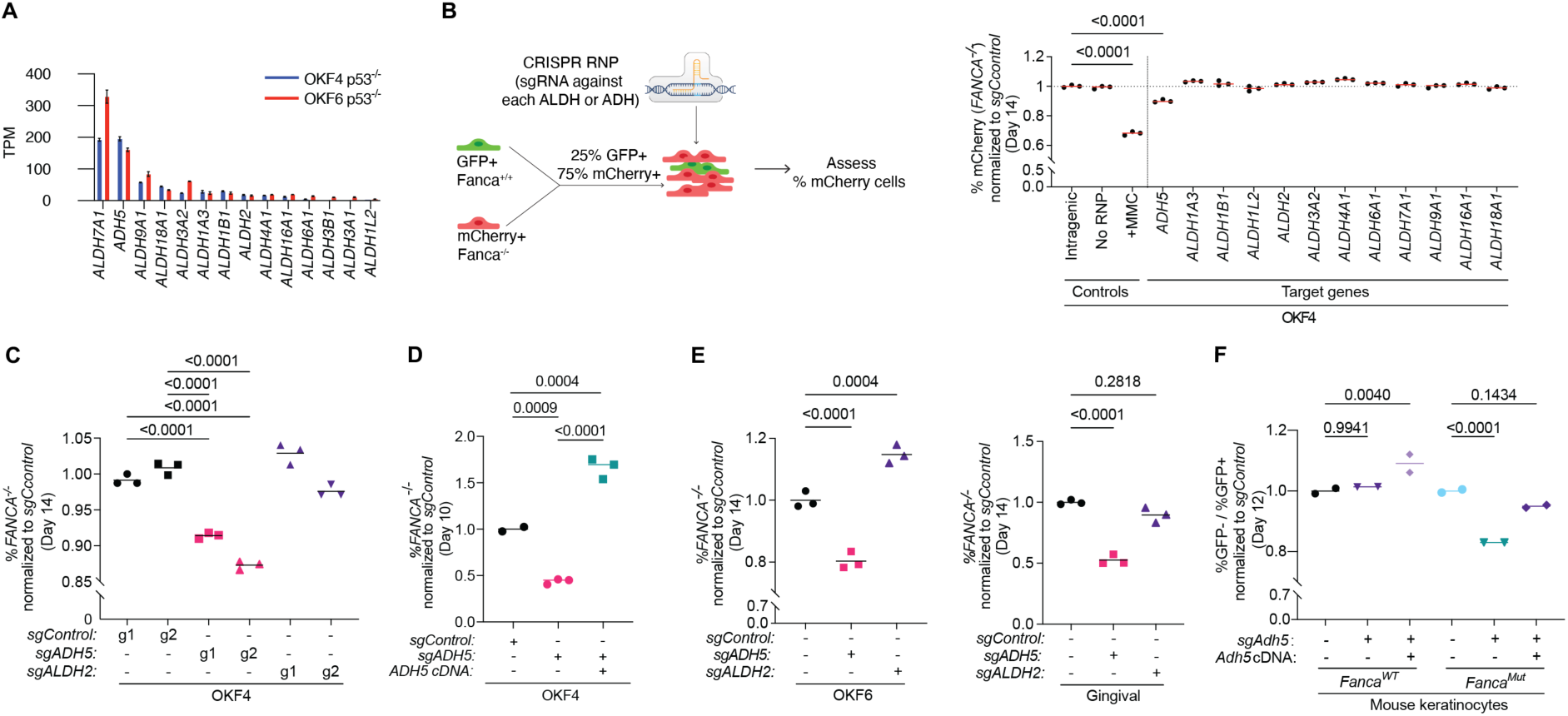
ADH5 knockout causes growth defects in FA-deficient human and mouse keratinocytes. (A) RNAseq expression of all *ADH* and *ALDH*-family genes with detectable expression (>2PTM) in OKF4 and OKF6 keratinocytes. (B) Competition assay in OKF4 with single-gene knockouts of *ADH* or *ALDH genes*. *Left*: Schematic of competition assay setup. *Right*: Quantification of competition assay after 14 days of culture. For each sample, the percentage of *FANCA^−/−^*(mCherry+) cells was normalized to the intragenic control. As a positive control, cells were treated with 25nM MMC for 4 days. The horizontal line indicates the mean of the intragenic control. (C) Competition assay with *ADH5* and *ALDH2* knockouts in OKF4 cells. For each sample, the percentage of *FANCA^−/−^* (mCherry+) cells was normalized to the sgControl cells. (D) Complementation experiment for *ADH5* knockout in OKF4 cells. Competition assay performed with ADH5 knockout complemented with empty vector (EV) and ADH5 cDNA. Relative abundance of *FANCA^−/−^* cells was quantified as described in (C). (E) Competition assays with *ADH5* and *ALDH2* knockouts in OKF6 (left) and gingival-Eh (right) keratinocytes. Relative abundance of *FANCA^−/−^* cells was quantified as described in (C). Synthego ICE analysis confirmed indel percentage of 95% (sgADH5) and 80% (sgALDH2) in OKF6, and 95% (sgADH5) and 88% (sgALDH2) in gingival-EH. (F) Competition assay in mouse keratinocytes with *Adh5* knockout FA-competent (*Fanca^WT^*) or FA-deficient (*Fanca^Mut^*) background. For each sample, the ratio of %GFP-over %GFP+ after 12 days of culture was normalized to the starting ratio at the beginning of experiment and further normalized to the *sgControl*. Synthego ICE analysis of indel percentages in mouse keratinocytes was 95% (sgADH5) in *Fanca^WT^*and 89% (sgADH5) in *Fanca^Mut^* cells. In B-F, each dot represents one technical replicate, and statistical significance was evaluated by one-way Kruskal-Wallis ANOVA.

With this information, we used ribonucleoprotein complex (RNP) delivery of CRISPR-Cas9 complexed with single sgRNAs to in-bulk cells knock out each of the 12 genes one by one and assess the effect with a competition assay in the OKF4 line, as schematized in **Figure 2B**. After two weeks of culture, only the knockout of *ADH5* led to a significant decrease in the percentage of FA-deficient cells (**Figure 2B and S2A**). The phenotype of ADH5 was validated using two independent guide RNAs targeting *ADH5* to control for off-target effects (**Figure 2C**) and further rescued through complementation of ADH5 (**Figure 2D and S2B and C**) in OKF4 cells. Notably, the complementation provided an additional growth advantage compared to FA-deficient cells expressing wild-type levels of ADH5. *ADH5* knockout also decreased cellular resistance to its aldehyde substrate, formaldehyde, and ADH5 complementation enhanced resistance compared to FA-deficient control cells (**Figure S2D**). We reasoned that the enhanced ability to metabolize endogenous and the exogenous formaldehyde accounts for the additional growth advantage observed in *ADH5*-complemented cells.

ADH5 is the principal enzyme responsible for formaldehyde detoxification in mammalian cells, with ALDH2 contributing in some contexts. OKF4 cells have low expression of *ALDH2* and its knockout did not show a significant drop in percentage of FA-deficient keratinocytes, suggesting that the keratinocytes depend mainly on ADH5 for formaldehyde detoxification (**Figure 2C**).

A similar dependence on ADH5 but not ALDH2 was also observed in OKF6 and gingival-EH keratinocytes (**Figure 2E**). In mouse keratinocytes, knockout of *Adh5* also induced synthetic lethality in FA-deficient cells, which was rescued by complementation of *Adh5* (**Figure 2F and S2E**).

Taken together, we demonstrate that ADH5 is involved in protection against endogenous formaldehyde in both human and mouse keratinocytes and its loss leads to decreased growth of FA-deficient cells.

### Loss of *ALDH3*-family genes leads to growth defects of FA-deficient human and mouse keratinocytes

ADHs and ALDHs are classified into families and subfamilies based on their shared amino acid sequence similarity and evolutionary relationship (*22*). Members of the same family are thought to have overlapping substrate specificities in addition to their distinct tissue distribution and efficiency. This redundancy may protect cells from reactive aldehydes and may explain why single-gene ALDH knockouts have minimal impact on aldehyde levels or growth of FA-deficient cells. Significant substrate accumulation typically requires simultaneous knockout of all family members.

Of the 19 human ALDH genes in 11 families, two families contain multiple members. One is the ALDH1/2 family with seven gene members that metabolize retinal, acetaldehyde and folate and the second is ALDH3-family with four gene members that metabolize fatty and medium/long chain aldehydes (*23, 24*). We focused on the ALDH3-family due to its high specificity for endogenous aldehydes from lipid peroxidation (LPO) and the LPO’s proposed role in skin diseases (reviewed in (*25*))

RNA-seq analysis of the four ALDH3-family genes (*ALDH3A1, ALDH3A2, ALDH3B1,* and *ALDH3B2*) revealed distinct expression patterns across keratinocyte cell lines (**Figure S1C**). Among the four isozymes, *ALDH3A2* is expressed in all four lines with high expression (>50 TPM) in OKF6 cells and moderate expression (10-50 TPM) in OKF4, gingival-EH, and mouse keratinocytes. In contrast, *ALDH3B1* and *ALDH3B2* exhibited low expression levels (1–10 TPM) across all keratinocyte lines except in OKF4 cells where there was no detectable expression (<1 TPM). *ALDH3A1* expression was only detected in OKF6 cells, but not in the other three lines. The differential expression of ALDH3-family genes across keratinocyte cell lines suggests that keratinocytes derived from different sites of the same tissue (e.g., the oral mucosa), may exhibit variable dependence on ALDH3-mediated defense mechanisms against LPO and LPO-derived metabolites.

In OKF6 cells, where all four *ALDH3* genes are expressed at a detectable level, we simultaneously knocked out *ALDH3A1, ALDH3A2, ALDH3B1,* and *ALDH3B2* using RNP delivery of CRISPR-Cas9 complexed with multiple sgRNAs. Efficient knockout of all four genes was confirmed by a combination of DNA sequencing of the targeted loci and immunoblotting (**Figure 3A**). The *ALDH3* quadruple knockout cells (sg*ALDH3^QKO^*) were sensitive to increasing dose of ML210, an inducer of lipid peroxidation (*26*), indicating that loss of the ALDH3-family diminishes protection against lipid peroxidation and related metabolites, including lipid aldehydes (**Figure 3B**). This increased sensitivity was rescued by expression of the *ALDH3A2* gene, chosen for its highest expression in keratinocytes. This is consistent with the functional redundancy among ALDH3-family enzymes in protection against lipid peroxidation-derived metabolites.

**Figure 3.**
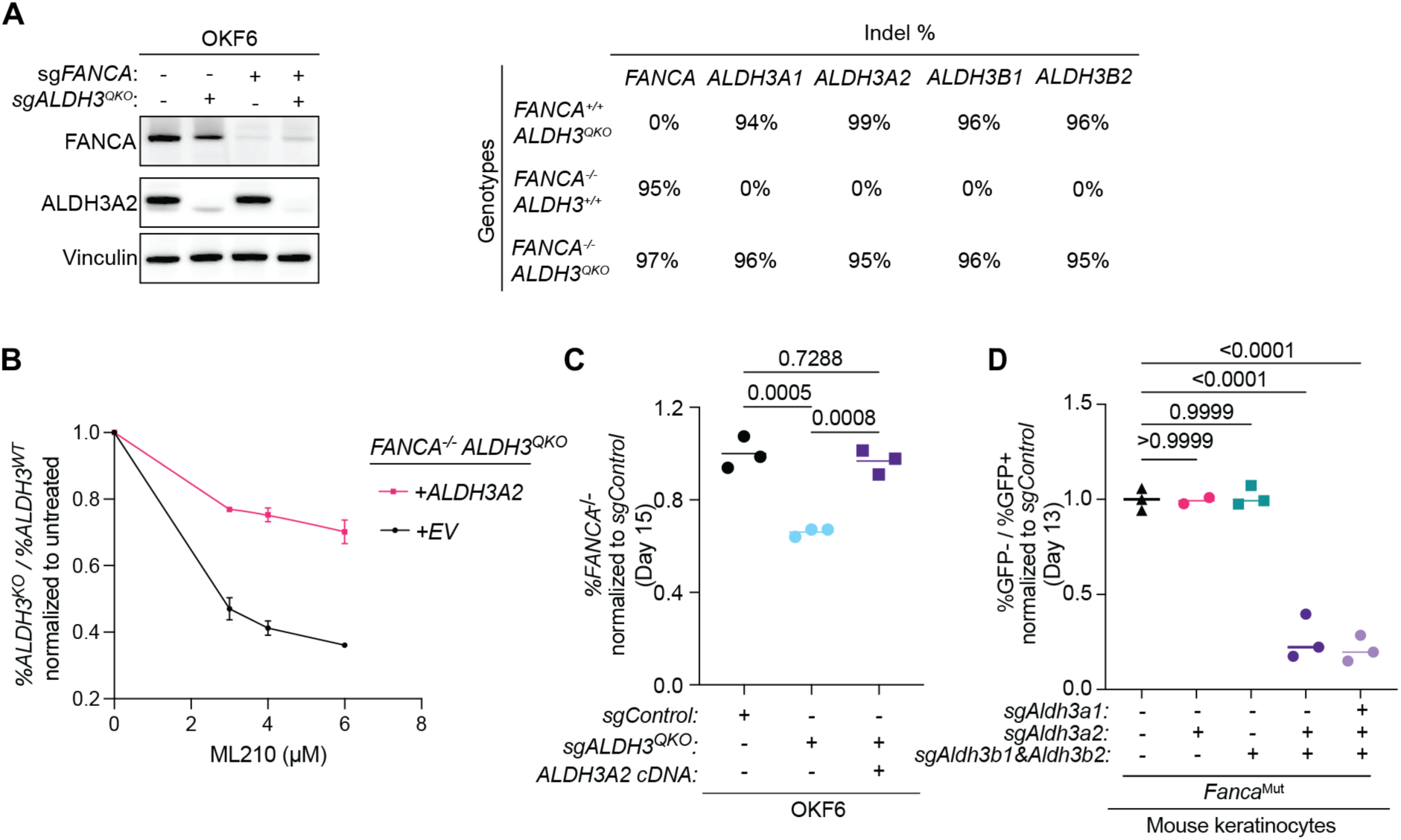
Loss of ALDH3 family genes leads to growth defects of FA-deficient human and mouse keratinocytes. (A) Validation of knockouts in OKF6 after *ALDH3-*family quadruple knockout*. Left*: Immunoblotting of FANCA and ALDH3A2 in OKF6 with *ALDH3* quadruple knockout (*ALDH3^QKO^*). Vinculin is used as loading control. *Right*: Summary of the percentage of insertion and deletion (indel) at each gene locus. (B) Sensitivity to the lipid peroxidation inducer ML210 in OKF6 *ALDH3^QKO^* knockout complemented with either empty vector (EV) or *ALDH3A2* cDNA, measured by competition assay. Cells were cultured with indicated dose of PFA and cultured for 3 days before they were analyzed by flow cytometry. The ratio of *FANCA^−/−^ ALDH3^QKO^* over *FANCA^−/−^* is calculated at each dose and normalized to the untreated control. Data are presented as mean ± SD. (C) Competition assay in OKF6 *ALDH3^QKO^* cells complemented with EV or *ALDH3A2* cDNA. (D) Competition assay in *Fanca^Mut^* mouse keratinocytes with *Aldh3a2* single knockout, *Aldh3b1* and *Aldh3b2* double knockout, *Aldh3* triple knockout and quadruple. For each sample, the ratio of %GFP-over %GFP+ cells after 16 days of culture was normalized to the starting ratio at the beginning of the experiment and then to sgControl. In C-D, each dot represents one technical replicate, and statistical significance was evaluated by one-way Kruskal-Wallis ANOVA.

The quadruple knockout of *ALDH3* in human cells led to a significant decrease in the percentage of FA-deficient cells compared to control, which could be rescued through complementation of *ALDH3A2* (**Figure 3C**). Mouse keratinocytes which express *Aldh3a2, Aldh3b1* and *Aldh3b2*, had no significant growth defect when only one or two of *Aldh3* genes were knocked out (*Aldh3a2* single knockout or *Aldh3b1*/*Aldh3b2* double knockout) in the setting of FA deficiency. In contrast, triple knockout of *Aldh3a2/Aldh3b1*/*Aldh3b2,* as well as quadruple knockout of all *Aldh3* enzymes led to a significant reduction in growth (**Figure 3D and S3A**). *ALDH3* quadruple knockout phenotype was rescued by complementation with *Fanca* cDNA (*Fanca^WT^)* and partially rescued by expression of *Aldh3a2* (**Figure S3B**).

In summary, we conclude that FA-deficient keratinocytes are more sensitive to *ALDH3-family* loss compared to their FA-competent counterparts in both human and mouse. Thus, we hypothesize that the growth of FA-deficient cells depends on efficient detoxification of LPO-derived aldehydes by the ALDH3-family members which are functionally redundant.

### Loss of ADH5 or ALDH3-family of enzymes induce DNA damage in keratinocytes

We reasoned that loss of ADH5 or ALDH3-family enzymes would lead to an accumulation of genotoxic aldehydes. These in turn would induce ICLs that would not be efficiently repaired in the setting of FA deficiency. In wild-type cells, ICLs block replication fork progression and are repaired by the coordinated action of FANC and accessory proteins. However, in the absence of FA pathway mediated repair, double-strand breaks (DSBs) form and persist, which we should be able to measure using phosphorylated histone variant H2AX (γH2AX) in cells that lacked ADH5 or ALDH3-family members.

Indeed, a combined knockout of *FANCA* and *ADH5* showed the highest γH2AX level in replicating (EdU+ positive) cells compared to control, *ADH5^−/−^* or *FANCA^−/−^*single knockout OKF4 cells (**Figure 4A**). This indicates that unrepaired DNA damage accumulates in FA-deficient keratinocytes when cells lose the ability to detoxify formaldehyde. In mouse keratinocytes, *Adh5*^−/−^ *Fanca*^−/−^ cells also exhibited the highest γH2AX level compared to control, *Adh5^−/−^*or *Fanca^−/−^* single knockouts. Complementation of *Adh5* cDNA effectively rescued the elevated γH2AX level, confirming that the increased DNA damage resulted from the loss of ADH5 (**Figure 4B**). Notably, *Adh5* knockout increased γH2AX levels in both FA-deficient and FA-competent backgrounds, indicating that formaldehyde detoxification by ADH5 is critical for maintaining genome stability even when the FA pathway is intact. This contrasts with findings in human OKF4s, where ADH5 loss alone induced relatively mild DNA damage, suggesting species- and cell line-specific differences in formaldehyde metabolism and DNA damage tolerance between mouse and human cells.

**Figure 4.**
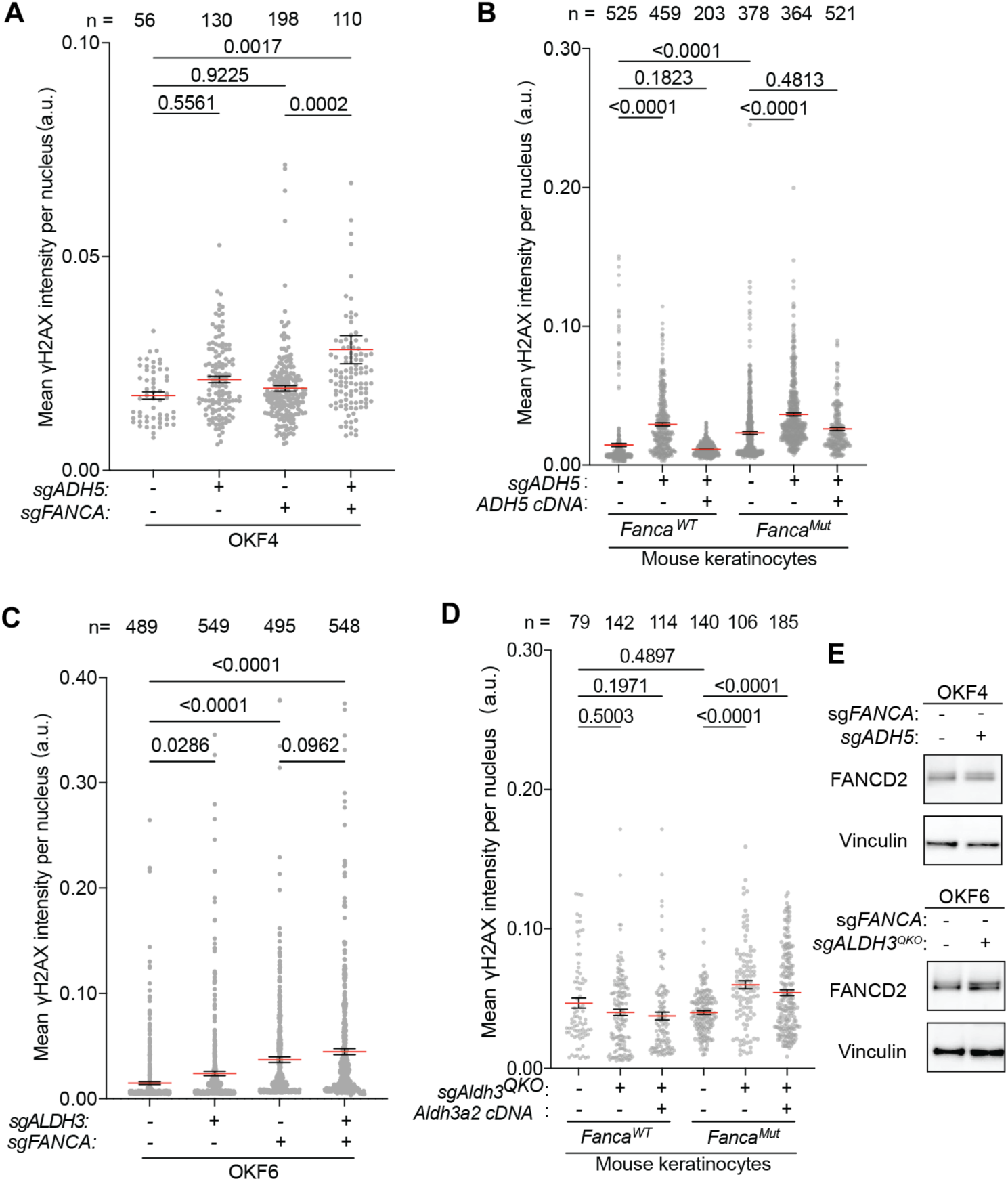
Loss of ADH5 or ALDH3 family enzymes induces DNA damage in keratinocytes. (A) Quantification of γH2AX in EdU+ *ADH5* knockout in OKF4 cells in FA-competent or FA-deficient backgrounds. EdU incorporation marks replicating S-phase cells. (B) Quantification of γH2AX in *Adh5* knockout mouse keratinocytes in FA-competent or FA-deficient backgrounds, complemented with empty vector (EV) and *Adh5* cDNA. (C) Quantification of γH2AX in EdU+ *ALDH3* quadruple knockout (*ALDH3^QKO^*) OKF6 cells in FA-competent or FA-deficient backgrounds. EdU incorporation marks replicating S-phase cells. (D) Quantification imaging of γH2AX in *Aldh3^QKO^* mouse keratinocytes in FA-competent or FA-deficient backgrounds, complemented with empty vector (EV) and *Aldh3a2* cDNA. (E) Immunoblotting of FANCD2 in *ADH5* knockout in OKF6 cells in FA-competent OKF4 and *ALDH3^QKO^* knockout in FA-competent. Vinculin is used as the loading controls. In A-D, each dot represents the mean intensity of γH2AX per nucleus. Data are presented as mean ± SEM. Statistical significance was evaluated by one-way Kruskal-Wallis ANOVA.

In OKF6 cells, a combined knockout of *FANCA* and all *ALDH3-*family genes exhibited the highest γH2AX level in EdU+ replicating cells compared to control, *ALDH3* quadruple knockout alone, or *FANCA^−/−^* single knockout alone (**Figure 4C**). We observed a similar phenotype in mouse keratinocytes, and complementation of *Aldh3a2* cDNA partially rescued the elevated γH2AX level (**Figure 4D**). Cell-cycle analysis showed significantly increased percentage of cells in late S/G2 phase in FA-deficient cells with *Aldh3* quadruple knockout, a phenotype seen in cells challenged with mild ICL exposure (**Figure S3C**). In summary, we propose that loss of ALDH3-family enzymes increase overall intracellular LPO-derived aldehyde burden that gives rise to DNA damage in FA-deficient keratinocytes, supporting the role of ALDH3-family enzymes in protection in both human and mouse keratinocytes.

Increased levels of γH2AX even in FA-proficient cells without ADH5 or ALDH3-family seen in the last experiments suggest that loss of the enzymes may be increasing the DNA damage. This is indeed the case, since FANCD2 ubiquitination was modestly increased upon knockout of *ADH5* or *ALDH3-family* genes (**Figure 4E**).

### FA-deficient keratinocytes are sensitive to treatment with formaldehyde, lipid peroxidation-inducing agents and lipid aldehydes

We have shown the importance of ADH5 and the ALDH3-family enzymes in protection of FA pathway-deficient keratinocytes. ADH5 is responsible for formaldehyde detoxification while ALDH3-family enzymes metabolize lipid aldehydes generated during lipid peroxidation. To determine if formaldehyde and LPO-derived aldehydes induce DNA damage in FA-deficient keratinocytes, we challenged FA-deficient and FA-competent mouse keratinocytes with paraformaldehyde (PFA), LPO-inducing ML210, and lipid aldehyde 4-hydroxynonenal (4HNE). As expected, FA-deficient cells were more sensitive to increasing doses of PFA, ML210 and 4HNE compared to FA-competent cells (**Figure 5A and S4C**). PFA, ML210 and 4HNE treatment greatly elevated γH2AX levels in FA-deficient cells and PFA and 4HNE slightly elevated γH2AX levels in FA-proficient cells (**Figure 5B**). The increase in DNA damage was accompanied by phosphorylation of p53 and upregulation of p21 (**Figure 5C**). Taken together, formaldehyde and LPO-derived aldehyde exposure causes DNA damage that activates the DNA damage response. Inability to repair DNA results in greater sensitivity of FA-deficient cells to aldehyde-induced DNA damage.

**Figure 5.**
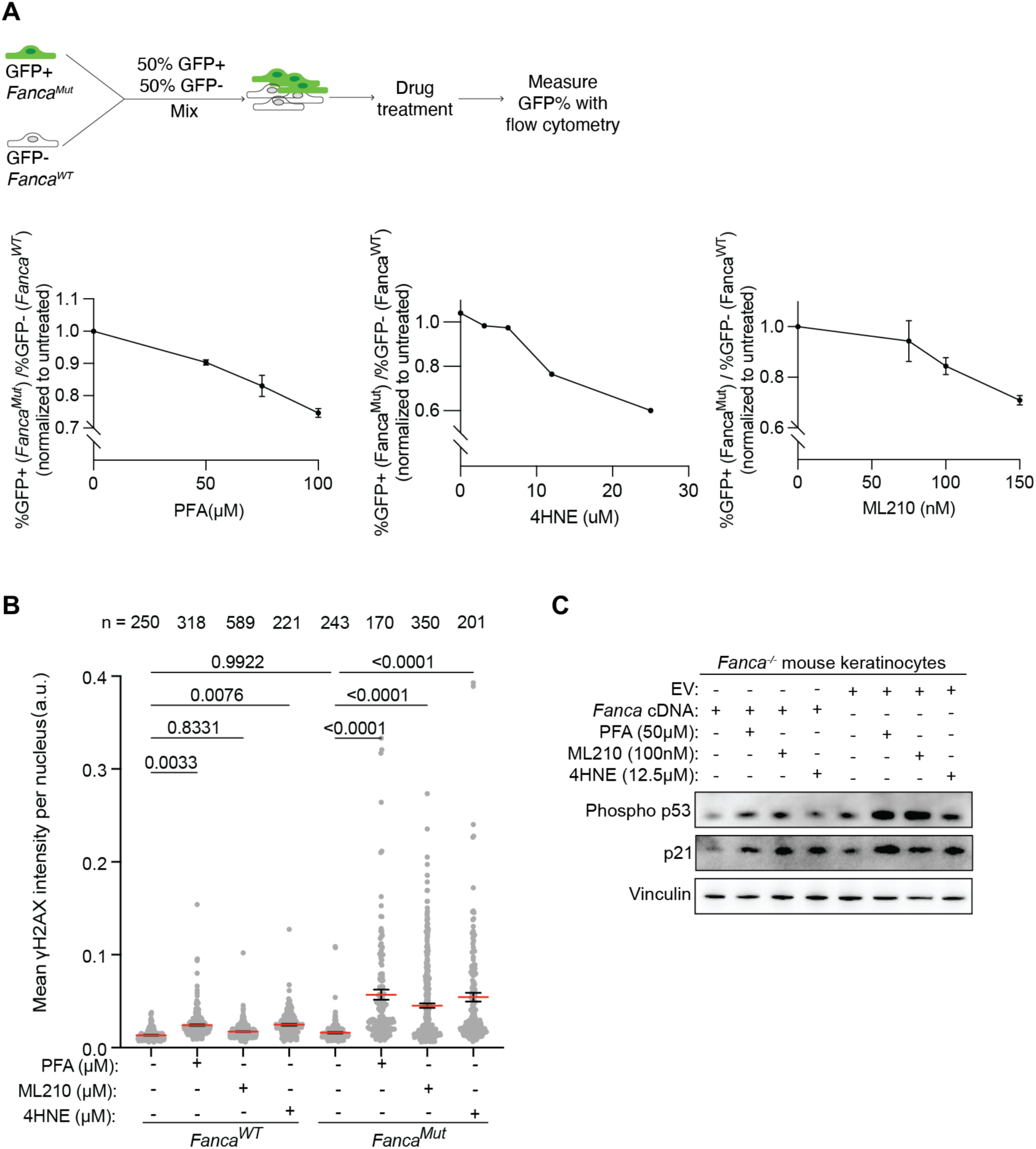
FA-deficient keratinocytes are more sensitive to formaldehyde, lipid peroxidation-inducing agent and lipid aldehydes than wild type keratinocytes. (A) Sensitivity to paraformaldehyde (PFA), ML210 and 4-hydroxynonenal (4HNE) in mouse keratinocytes, measured by competition assay. Cells were treated with paraformaldehyde (PFA), ML210 and 4-hydroxynonenal (4HNE) for 24h. *Top*: Schematic of the competition assay setup. *Bottom*: Quantification of competition assay. For each condition, the ratio of %GFP-over %GFP+ cells was calculated and normalized to the untreated control. Data are presented as mean ± SD. (B) Quantification of γH2AX in FA-competent (*Fanca^WT^*) and FA-deficient (*Fanca^Mut^*) mouse keratinocytes treated with PFA, ML210 and 4HNE for 24h. Each dot represents the mean γH2AX intensity per nucleus. Data are presented as mean ± SEM. Statistical significance was evaluated by one-way Kruskal-Wallis ANOVA. (C) Immunoblotting of phospho-p53 and p21 in FA-competent (*Fanca^WT^*) and FA-deficient (*Fanca^Mut^*) mouse keratinocytes following treatment of PFA, ML210 and 4HNE. Vinculin is used as a loading control.

### Treatment with the antioxidant N-acetylcysteine rescues growth of FA-deficient ADH5 knockout keratinocytes

Our lab has previously demonstrated that FA-associated SCCs are characterized by high levels of structural variants leading to copy number alterations in oncogenes and tumor suppressor genes, and have an aggressive course with limited therapeutic options (*10*). Prevention of mucosal SCCs is a priority in this patient population. Having identified formaldehyde as an important aldehyde in mucosal keratinocytes, we explored compounds that are known to scavenge aldehydes. Thiol-rich antioxidant N-acetyl-L-cysteine (NAC) has been shown to abate formaldehyde toxicity and has been proposed to scavenge formaldehyde by either directly reacting with formaldehyde or replenishing the pool of intracellular GSH (*27*). We also tested metformin (*N*,*N*-dimethylbiguanide), a widely used drug to treat diabetes with proven safety. As a guanine derivative, metformin has the potential to scavenge DNA damage-inducing aldehydes through the Mannich reaction. Metformin has been shown to protect FA-deficient hematopoietic stem cells from DNA damage and delay onset of tumors in tumor-prone FA-deficient *Fancd2^−/−^ Trp53^+/-^* mice. It was safe and tolerable in nondiabetic patients with FA and was deemed to have potential to provide therapeutic benefit for FA patients, although human studies have been limited (*28, 29*).

To test if supplementation of growth media with NAC or metformin rescues the growth of *FANCA^−/−^ ADH5^−/−^* we once again turned to a competition assay (**Figure 6A**). Cells were treated daily with 250μM NAC or metformin, a relatively low dosage where *FANCA^−/−^* cell growth was not affected by the drug treatment (**Figure S5B**). After 12-16 days of treatment, the percent of *FANCA^−/−^*

**Figure 6.**
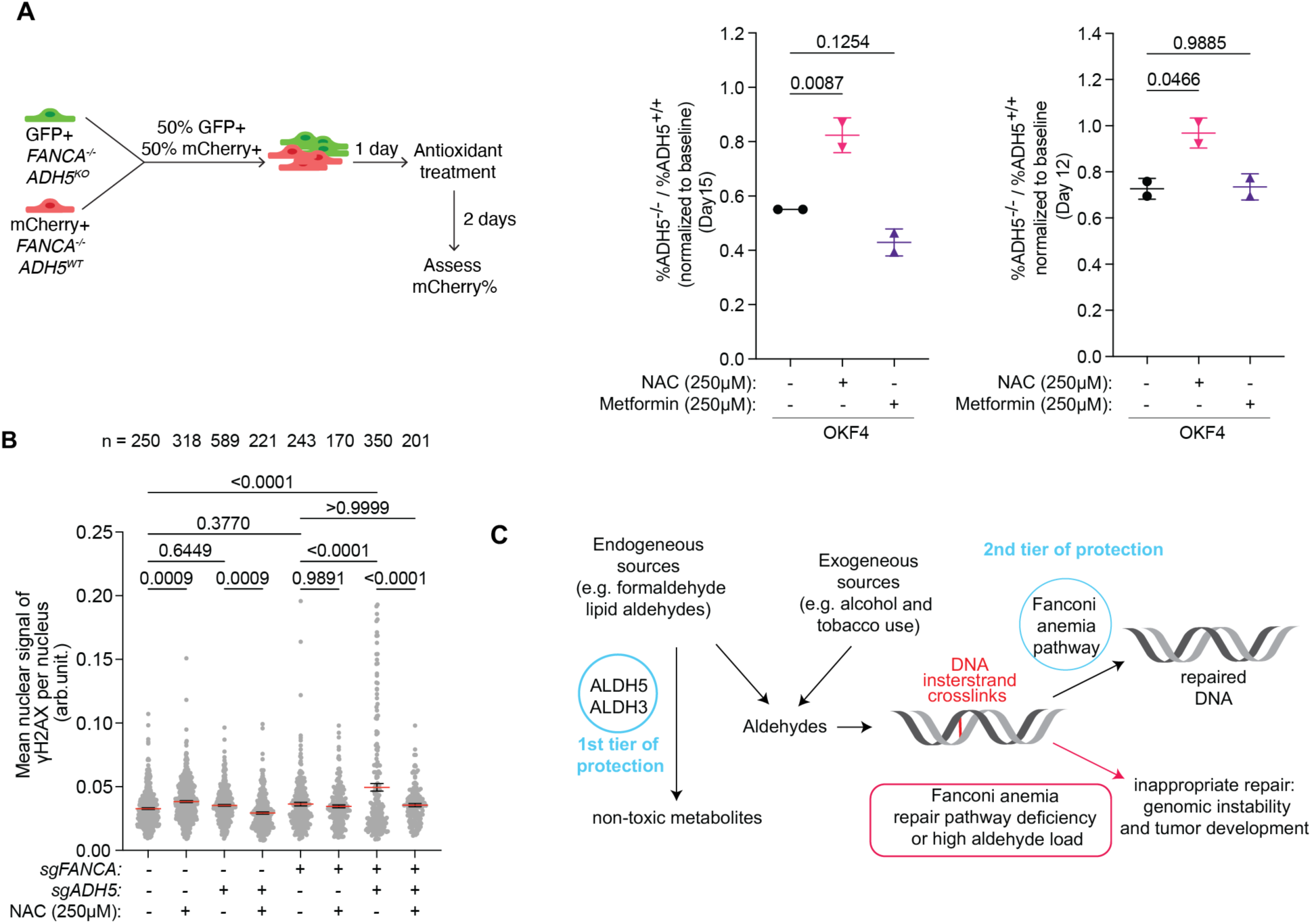
Antioxidant N-acetylcysteine rescues the growth of FA-deficient ADH5 knockout keratinocytes. (A) Competition assay in co-cultured *FANCA^−/−^* and *FANCA^−/−^ ADH5^−/−^* OKF4 cells treated daily with 250μM N-acetylcysteine (NAC) or 250μM metformin. The ratio of % *FANCA^−/−^ ADH5^−/−^* over % *FANCA^−/−^* cells at the end of the experiment was normalized to the starting ratio at the beginning of experiment. Each dot represents one technical replicate, and statistical significance was evaluated by one-way Kruskal-Wallis ANOVA. (B) Quantification of γH2AX in *ADH5* knockout OKF4 cells in FA-competent or FA-deficient backgrounds, with and without daily 250μM NAC treatment. Each dot represents the mean γH2AX intensity per nucleus. Data are presented as mean ± SEM. Statistical significance was evaluated by one-way Kruskal-Wallis ANOVA. (C) Schematic illustrating the two-tier protection mechanism against formaldehyde and lipid aldehydes in keratinocytes. Tier 1: Enzymatic detoxification of formaldehyde and lipid aldehydes. Tier 2: DNA interstrand crosslink repair by the Fanconi anemia pathway. FA-deficient keratinocytes that lack the tier 2 protection rely on detoxification of formaldehyde and lipid aldehydes (tier 1) to prevent genome instability that contributes to tumorigenesis.

*ADH5^−/−^* cells relative to *FANCA^−/−^* cells was significantly rescued by NAC treatment compared to the untreated (**Figure 6A and S5A**). NAC treatment also significantly lowered γH2AX level in *FANCA^−/−^ ADH5^−/−^* cells, as well as *FANCA^+/+^ ADH5^−/−^* cells, confirming that NAC can decrease markers of DNA damage in both FA-proficient and FA-deficient keratinocytes that have lost ADH5 (**Figure 6B**). Metformin treatment did not show significant rescue, indicating that it may not act as a direct scavenger of formaldehyde.

## Discussion

Using a sensitized cellular system deficient in the Fanconi anemia pathway, we demonstrated that ADH5 and the ALDH3-family of enzymes protect mammalian keratinocytes from endogenous DNA damage. These enzymes, which detoxify formaldehyde and lipid-derived aldehydes, respectively, constitute the first tier of defense against DNA-damaging metabolites that can mutagenize oral and other squamous mucosa stem cells and set the stage for cancer development (**Figure 6C**). The FA pathway acts as a second tier of defense to remove the ICLs induced by aldehydes that escaped the detoxification by ADH5 and ALDH3-family enzymes. We posit that even a low level of those escaped aldehydes, potentially in combination with exogenous aldehydes found in food, alcohol, tobacco, and polluted air, could lead to mucosal SCCs in patients with FA, who lack the second tier of defense against ICLs (*30–34*). However, the causal relationship between the loss of ADH5 or ALDH3*-*family genes and FA-associated SCC onset remains to be validated *in vivo*. Ongoing efforts are focused on developing mouse models that spontaneously develop oral SCCs, which will enable mechanistic studies to confirm the protective roles of ADH5 and ALDH3 enzymes in maintaining genome stability and preventing cancer development. Such models will also provide a valuable platform for evaluating aldehyde-scavenging and other compounds as potential cancer-prevention strategies.

Loss or insufficient activity of ALDH2 and ADH5, which both metabolize formaldehyde, has been linked to early bone marrow failure and the spontaneous development of leukemia in mice and humans with FA pathway deficiency, resulting in a DNA repair defect. Similar clinical phenotypes are also seen in patients with Aldehyde Degradation Deficiency Syndrome (combination of *ALDH2* and *ADH5* deficiency), who have normal DNA repair but increased formaldehyde load (*13–19*). Based on our findings that ADH5 protects keratinocyte genomes, we anticipate that individuals with Aldehyde Degradation Deficiency Syndrome may be at increased risk for mucosal squamous cell carcinomas. Although oral cancers have not yet been reported in these patients, likely because all identified individuals are under the age of 19, screening for oral, esophageal, and anogenital cancers would be prudent.

Knockout of *ALDH2* in Fanconi anemia pathway-deficient keratinocytes did not have a significant synthetic lethal effect in our study. It is possible that ALDH2 still contributes to genome protection in keratinocytes, but its role may be masked by the dominant activity of ADH5. Unlike in the bone marrow, this redundancy was not immediately apparent in keratinocytes. However, simultaneous loss of ALDH2 and ADH5 could uncover such functional overlap, even in keratinocytes.

In this study, we also uncovered functional redundancy in keratinocyte genome protection within the ALDH3-family of the ALDH3A1, ALDH3A2, ALDH3B1, and ALDH3B2 enzymes. In the keratinocyte lines used, the most abundant family member of ALDH3-family is *ALDH3A2*, which oxidizes medium- and long-chain aliphatic aldehydes to fatty acids. In human tissues, ALDH3A2 is broadly expressed, with highest expression in liver, gallbladder, endocrine tissues, skin, and genitourinary tract (*35, 36*). ALDH3A2 mutations cause lipid accumulation in skin and brain membranes, leading to Sjögren-Larsson syndrome (*37*). ALDH3B1 metabolizes medium- and long-chain fatty aldehydes and provides protection from 4-HNE toxicity (*38*). It is also broadly expressed, with higher levels in bone marrow, lymphoid tissues, fallopian tubes, and respiratory epithelium (*35, 36*). The two remaining ALDH3-family members are ALDH3A1 and ALDH3B2, both of which show very low or absent expression in most human tissues (*35, 36*). ALDH3A1 primarily metabolizes medium-chain saturated and unsaturated aldehydes, such as 4-HNE. It shows moderate to high expression in the digestive tract and low expression in skin, however its high expression in corneal epithelial and stromal cells is particularly important for protection of the cornea (*39–41*). Lastly, ALDH3B2, along with the rodent-specific ALDH3B3 (orthologous to human ALDH3B1), exhibits broad substrate specificity for similar aldehydes. It shows limited expression overall but higher levels in the proximal digestive tract, skin, and breast tissue (*35, 36*).

Taken together, there is significant functional redundancy among the enzymes of the ALDH3 family. All four family members seem to be involved in tier 1 defense mechanism in keratinocytes, since removal of all four was needed to elicit synthetic lethality in combination with FA deficiency. However, we still have not identified the most deleterious of the lipid aldehydes in keratinocytes.

There are many potential sources of lipid aldehydes in keratinocytes including lipid peroxidation (LPO) of polyunsaturated fatty acids (PUFAs) within membrane phospholipids, typically triggered by reactive oxygen species. Multiple studies have demonstrated that lipid peroxidation can also affect nuclear membranes, which could lead to the generation of such lipid aldehydes in the nucleus (*42, 43*). Keratinocytes undergo dramatic remodeling of lipid metabolism as they differentiate which may also create DNA damaging metabolites. Undifferentiated keratinocytes have higher proportion PUFAs and monounsaturated fatty acids that are replaced by saturated fatty acids and very-long-chain fatty acids upon differentiation (*44*).

Lipid composition, particularly PUFA content, in keratinocytes has not been well characterized in the context of Fanconi anemia. Emerging evidence suggests that oral keratinocytes display altered lipid dependencies upon loss of the FA-pathway, including upregulation of multiple specific of glycosphingolipids such as the GM3 ganglioside, which promote invasive behavior (*45*). ALDH3-family members catalyze the oxidation of long-chain fatty aldehydes including C16 aldehydes (hexadecanal and trans-2-hexadecenal) generated through sphingolipid metabolism (*46*). It remains to be seen if sphingolipids or the increased levels of sphingolipids are the cancer-driving sources of ICL-inducing metabolites besides endogenous formaldehyde.

A remaining challenge is to decrease the genotoxic burden arising from endogenous formaldehyde and lipid aldehydes in mucosal keratinocytes. We show that NAC lowered DNA damage and improved growth of *ADH5^−/−^* keratinocytes, in both an FA-competent and FA-deficient setting, confirming the potential of aldehyde scavengers, particularly for formaldehyde and LPO-derived aldehydes, as cancer prevention strategy. We believe that better understanding of endogenous aldehyde sources will explain the high predisposition of patients with FA to early onset mucosal cancers and identify potential preventive strategies for patients with FA and those with sporadic cancers.

## Materials and Methods

### Keratinocyte cell line generation and culture

The Gingival-EH line was derived from primary Gingival Keratinocytes (PCS-200-014, ATCC). Primary keratinocytes were immortalized with pWZL-MSCV-hTERT and MSCV-based HPV E6E7 expression with retroviruses produced 293T cells (ATCC CRL-3216) as described below. The OKF4 and OKF6 cell lines were obtained from Dr. Cathie Garnis (originally generated by Dr. James Rheinwald with constitutive hTERT expression to establish the cell line) (*47, 48*). To immortalize these lines, we generated a bulk knockout of *TP53* using CRISPR Cas9 ribonucleoprotein delivery (RNP, IDT). sgRNAs sequences are listed in **Table S1.** Immortalized Gingival-EH, OKF4 and OKF6 lines were cultured in KGM-gold keratinocyte media supplemented with 50µM Calcium Chloride and Bulletkit growth supplements (Lonza Biosciences). Mouse keratinocyte lines were generated as described before (*10*). All keratinocytes were cultured at ambient CO_2_/3% O_2_ at 37 °C. For passaging, cells were dissociated with TrypLE. After the reaction was quenched with 2% calcium-chelated FBS, cells were replated.

### Viral transductions

cDNAs and CRISPR Cas9 vectors were delivered by retroviral or lentiviral transduction after packaging in HEK293T cells. To package the viruses, 5 × 10^6^ 293T cells were plated the day before transfection. DNA and viral packaging vectors were transfected into cells using TransIT 293 transfection reagent (Mirus Bio) according to the manufacturer’s protocol. Media was changed the next day. Viral supernatant was harvested and filtered (0.45 μM) at 24 and 48 hrs post-transfection and then incubated with LentiX Concentrator (Takata Bioscience) according to the manufacturer’s protocol to be concentrated and resuspended in appropriate keratinocyte media. Target cells were infected with viral supernatants supplemented with 4 μg/mL polybrene. Media was changed the following day. Two days after infection, stably expressing cells were selected with the appropriate antibiotics. The packaging vectors pMD2.G (Addgene plasmid # 12259) and psPAX2 (Addgene plasmid # 12260) for lentiviruses were gifted by Didier Trono. VSV-G and Gagpol were used for retroviral packaging. All plasmids are listed in **Table S2**.

### Generation of FA-deficient and FA-competent human and mouse keratinocyte cell lines

To knock out FANCA in human keratinocytes, we delivered CRISPR Cas9 and sgRNAs against human *FANCA* gene (*FANCA^−/−^*) or a random intragenic region (control, *FANCA^+/+^)* through either lentiCRISPRv2 lentiviral transduction (Plasmid #52961, Addgene) or ribonucleoprotein complex (RNP) Alt-R CRISPR-Cas9 system (IDT). For competition assays in Figure 2B and C, cells were infected with lentiCRISPRv2-sgFANCA-Cas9-IRES-mCherry or LCV2-sgControl-IRES-GFP lentiviruses. Guides were cloned into LentiCRIPSR v2 (LCV2) plasmid with Golden Gate cloning according to previous protocol (*49*)with modifications including: (1) LCV2 vector was used instead of pX330 vector, (2) BsmBI-v2 was used instead of BbsI and (3) CutSmart buffer was used. Cells were subsequently sorted by FACS for mCherry^+^ and GFP^+^ for FA-deficient and FA-competent cells, respectively. For all the other experiments, *FANCA* knockout was achieved using RNP following the manufacturer’s protocol. Cells were then labeled with GFP or mCherry using pLenti-GFP or pLenti-mCherry lentiviruses and were FACS-sorted.

For the mouse keratinocytes, *Fanca^−/−^* cells were isolated and immortalized from neonate skin. FA-competent cells (*Fanca^+/+^*) were generated through complementation with pMSCVhyg-*Fanca* retrovirus. The pMSCVhyg-*Fanca* plasmid was generated through gateway cloning. Finally, *Fanca^−/−^* cells were labeled with GFP using pLenti-GFP lentivirus and were FACS-sorted.

### MMC sensitivity assay

70,000 cells were plated per well of 6-well plates. The following day, cells were treated with the indicated doses of Mitomycin-C (M4287, Millipore Sigma) in triplicate. 4 days later, cells were counted using a Z2 Coulter Counter (Beckman Coulter). The number of surviving cells was normalized to untreated controls.

### Fluorescence-based competition assay and flow cytometry

Competition assays were used to measure the growth of FA-deficient keratinocytes relative to their FA-competent pair upon knockout of an *ADH* or *ALDH* gene(s). For human keratinocytes, mCherry^+^ cells with one genotype were mixed with GFP^+^ cells of a second genotype at an appropriate ratio (see schematics in the Figures) and were plated at 50,000-80,000 cells per well in 6-well plates. Cells were then passaged every 7 days, and a portion of the cell mixture was harvested for flow cytometry analysis at each time point. The ratio of mCherry^+^ cells and GFP^+^ cells was obtained at the beginning of the experiment (baseline) and at each time point. At the end of the experiment, the ratio was first normalized to control. P-values were calculated with one-way ANOVA. For mouse keratinocytes, GFP^−^ cells with indicated ADH5 or ALDH3 knockout were mixed with GFP^+^ control *Fanca^+/+^* or *Fanca^−/−^* cells. 70,000 cells were plated per well in 6-well plates and passaged every 3 to 4 days and analyzed as in experiments with human keratinocytes.

For testing hypersensitive to pharmacological perturbations, cells were plated as described above and treated with indicated doses of MMC, paraformaldehyde (#15710, EMS), ML210 (HY-100003, MedChemExpress) or 4-hydroxynonenal (#32100, Cayman Chemical) on the following day. The media remained unchanged until 4 days later when cells were harvested for flow cytometry analysis. For human keratinocytes, the ratio of mCherry^+^ cells and GFP^+^ cells was calculated at each concentration and normalized to the untreated sample. For mouse keratinocytes, the ratio of GFP^−/^GFP^+^ was calculated at each concentration and normalized to the untreated. P-values were calculated with one-way ANOVA.

To prepare harvested cells for flow cytometry analysis, cells were fixed in 100µL 4% PFA and permeabilized with ice-cold 70% ethanol. The samples were stored at −20°C until analysis on the NovoCyte Penteon flow cytometer (Agilent). Prior to analysis, samples were washed with PBS twice and resuspended in PBS. Data were visualized and analyzed with FlowJo.

### RNA Sequencing (RNA-Seq)

Total messenger RNA was extracted using RNeasy Plus Mini kits (Qiagen). Samples were sent to Azenta-Genewiz for RNA purity analysis, mRNA selection (by PolyA selection), cDNA library preparation, and paired-end sequencing at a depth of >30 million reads per sample (Illumina HiSeq, 2×150bp, single index). RNA Sequencing analysis was performed by the Bioinformatics core at the Rockefeller University using the NGSpipeR and Salmon packages, and gene-level normalized transcript per millions (TPM) values are shown in **Table S3.**

### CRIPSR Cas9 knockout of ALDHs and ADHs and validation of knockout and complementation

For experiments shown in Figure 2B, each ALDH or ADH gene was knocked out through RNP delivery of CRISPR Cas9 and one sgRNA targeting the gene. For knockouts of ADH5 in all other experiments, we utilized RNP to deliver CRISPR Cas9 and a single sgRNA targeting human *ADH5* or mouse *Adh5* gene. The knockout efficiency was validated through PCR of the target gene locus from genomic DNA isolated from the knockout cells with overnight cell lysis using DirectPCR buffer (Viagen) containing 1:50 Proteinase K. The PCR product was sequenced with Sanger sequencing, and the percentage of insertion and deletion was calculated with EditCo ICE analysis tool. Functional loss of ADH5 was tested as hypersensitivity to paraformaldehyde (PFA) treatment in competition assay. Once knockout was validated, cells were complemented with empty vector (EV) or cDNA encoding for human or mouse ADH5 through infection with pMSCVneo-*EV* or pMSCVneo-*ADH5 cDNA* retroviruses, respectively. The pMSCVneo-*EV* or pMSCVneo-*ADH5 cDNA* plasmids were generated through Gateway cloning.

For the quadruple knockout of ALDH3, RNP was used to deliver CRISPR Cas9 and three sgRNAs simultaneously. *ALDH3A1* and *ALDH3A2* genes were each targeted by one sgRNA, whereas another sgRNA targeted a shared sequence between *ALDH3B1* and *ALDH3B2.* The knockout efficiency was validated through sequencing of each target gene locus, and percentage of insert and deletion was calculated with EditCo ICE analysis tool. Loss of ALDH3A2 protein expression was tested using immunoblotting. Functional loss of ALDH3-family enzymes was tested as hypersensitivity to ML210 treatment in competition assay. Once knockout was validated, cells were complemented with empty vector (EV) or cDNA encoding the human or mouse ALDH3A2 through infection with pMSCVneo-*EV* or pMSCVneo-*ALDH3a2 cDNA* retroviruses, respectively.

### Immunofluorescence staining

Cells were plated on cover slips in 24-well plates. After seeding, cells were treated with indicated compound or were pulsed with 10μM EdU for 1.5 hours. After washing with PBS once, cells were permeabilized with 0.5% Triton X-100 (Sigma-Aldrich) in PBS for 5 min on ice. Cells were then washed twice with PBS and fixed with 3.7% formaldehyde (Sigma-Aldrich) in PBS for 10 min at room temperature. EdU was stained with the Click-iT^TM^ EdU Alexa Fluor^TM^ 647 imaging kit (Invitrogen) according to manufacturer’s protocol. After washing with PBS twice, cells were incubated in 3% BSA in PBS for 1 h at room temperature for blocking, followed by incubation with primary antibody at 4°C overnight. After washing cells three times with PBS, the secondary antibody was incubated for 1 h at room temperature. Then, cells were washed three times with PBS and incubated with 1μg/ml DAPI solution in PBS. Finally, the cover slips were mounted with Fluoromount-G (Invitrogen).

Panels of images were acquired on an inverted Olympus IX-71 DeltaVision Image Restoration Microscope (Applied Precision) using a 20X objective. CellProfiler was used to generate a nuclear mask based on DAPI signals. The mean intensities of γH2AX and EdU for each nucleus were then measured within the nuclear mask. Output data from CellProfiler was visualized in GraphPad Prism.

### Western blotting

Whole cell lysates were prepared by lysing cells in RIPA buffer (0.1% SDS, 1% Triton X-100, 1% sodium deoxycholate, 1mM EDTA, 150mM NaCl, 50mM Tris-HCl (pH=6.8)) and protein concentration was determined using the DC protein assay (Bio-Rad) prior to addition of 2-mercaptoethanol and bromophenol blue. DNA was sheared by sonication and samples were heated at 95C for 10 minutes. Proteins were separated by sodium dodecyl sulfate polyacrylamide gel electrophoresis (SDS-PAGE) on precast 4–12% Bis-Tris or 3-8% Tris-Acetate gels. Membranes were blocked for 1hr in 5% milk in TBST (10 mM Tris-HCl (pH=7.5), 150 mM NaCl, 0.1% Tween 20) and incubated in primary antibodies overnight at 4 °C. Membranes were washed in TBST (3 × 10 minutes) then incubated with horseradish peroxidase (HRP)-conjugated secondary antibodies for 1hr at room temperature. Membranes were washed in TBST (3 ×10 minutes) and detected by enhanced chemiluminescence using an Azure c300 imaging system.

Antibodies included FANCA (Bethyl Labs, A301-980A, 1:1000), FANCD2 (Novus Biologicals, NB100-182, 1:1000), Vinculin (MilliporeSigma, V9131, 1:5000), ALDH3A2(Sigma, SAB3500221, 1:1000), PhosphoP53 (Cell Signaling Technology, #9284, 1:1000), p21 (Santa Cruz, sc-6246, 1:200) and γH2AX (Millipore Sigma, 05-636, 1:1000).

### Treatment of antioxidants

N-acetyl-L-cysteine (Sigma) and metformin hydrochloride (Sigma) were resuspended in KGM-gold media as 200mM and 150mM stock, respectively. 7,000 OKF4 cells were plated per well in 12-well plates. After seeding, the stock solution of NAC was diluted to 250μM in KGM-gold media for treatment. The media was refreshed daily.

### Statistics

Significance was evaluated either by Kruskal-Wallis ANOVA with a Dunn’s post-test or by unpaired parametric t test using GraphPad Prism software. All p values are shown in the figure panels, and which test was used is indicated in the figure legend. Descriptions of the statistical analyses presented in the figures can be found in the corresponding figure legends.

## Acknowledgments

We are grateful to all the members of the Smogorzewska laboratories for helpful discussions throughout the project. We thank Dr. Meng Wang for experimental advice. We thank Dr. Cathie Garnis and Dr. James Rheinwald for providing us with the OKF4 and OKF6 cell lines and Dr. Markus Grompe for providing *Fanca* mutant mice. We gratefully acknowledge the Bioinformatics Resource Center, the Bio-Imaging Resource Center, the Flow Cytometry Resource Center, and the Genomics Resource Center at the Rockefeller University for experimental support.

## Author contributions

Y.Y., N.J.B, and A.S. designed research; Y.Y., N.J.B performed research; Y.Y., N.J.B. analyzed data; and Y.Y., N.J.B. and A.S. wrote the paper.

## Supplemental Data

**Supplemental Figure 1.**
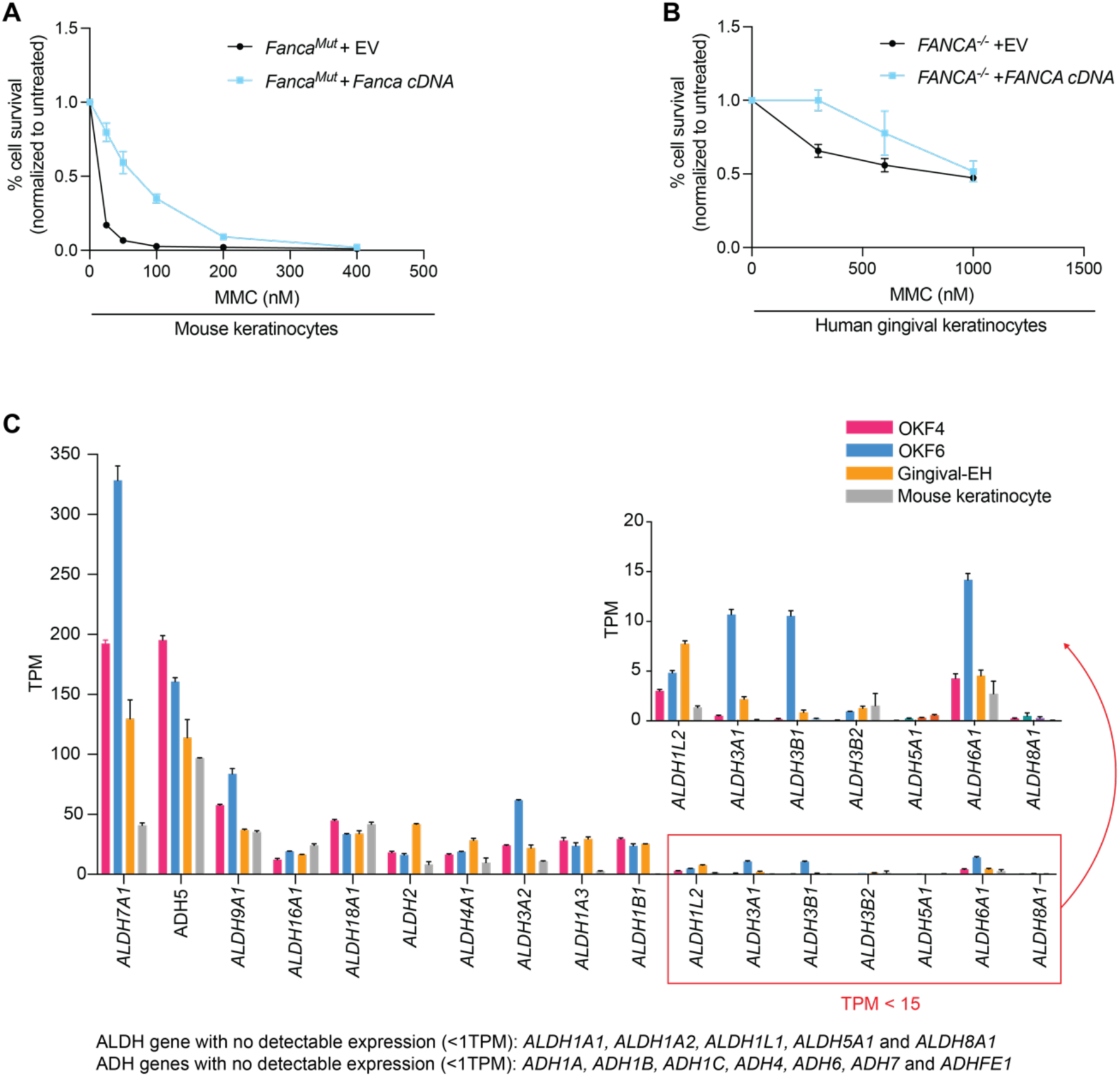
Further characterization of keratinocyte cell lines from human and mouse. (A) MMC sensitivity assay in mouse keratinocytes. FA-deficient (*Fanca^Mut^*) keratinocytes were complemented with empty vector (EV) or *Fanca* cDNA. Cells were treated with indicated dose of MMC and then cultured for 5 days before cells were counted to calculate percent of cell survival. (B) MMC sensitivity assay in human *FANCA^−/−^* gingival-Eh cells complemented with EV and *FANCA* cDNA. Experiment was carried out as in (A). (C) RNAseq expression levels of all *ADH* and *ALDH* family genes in the OKF4, OKF6, gingival-Eh, and mouse keratinocyte lines. On the top right, a zoomed-in plot shows genes with low expression (TPM<15). Genes with no detectable expression (TPM<1) are summarized at the bottom. See Table S3.

**Supplemental Figure 2.**
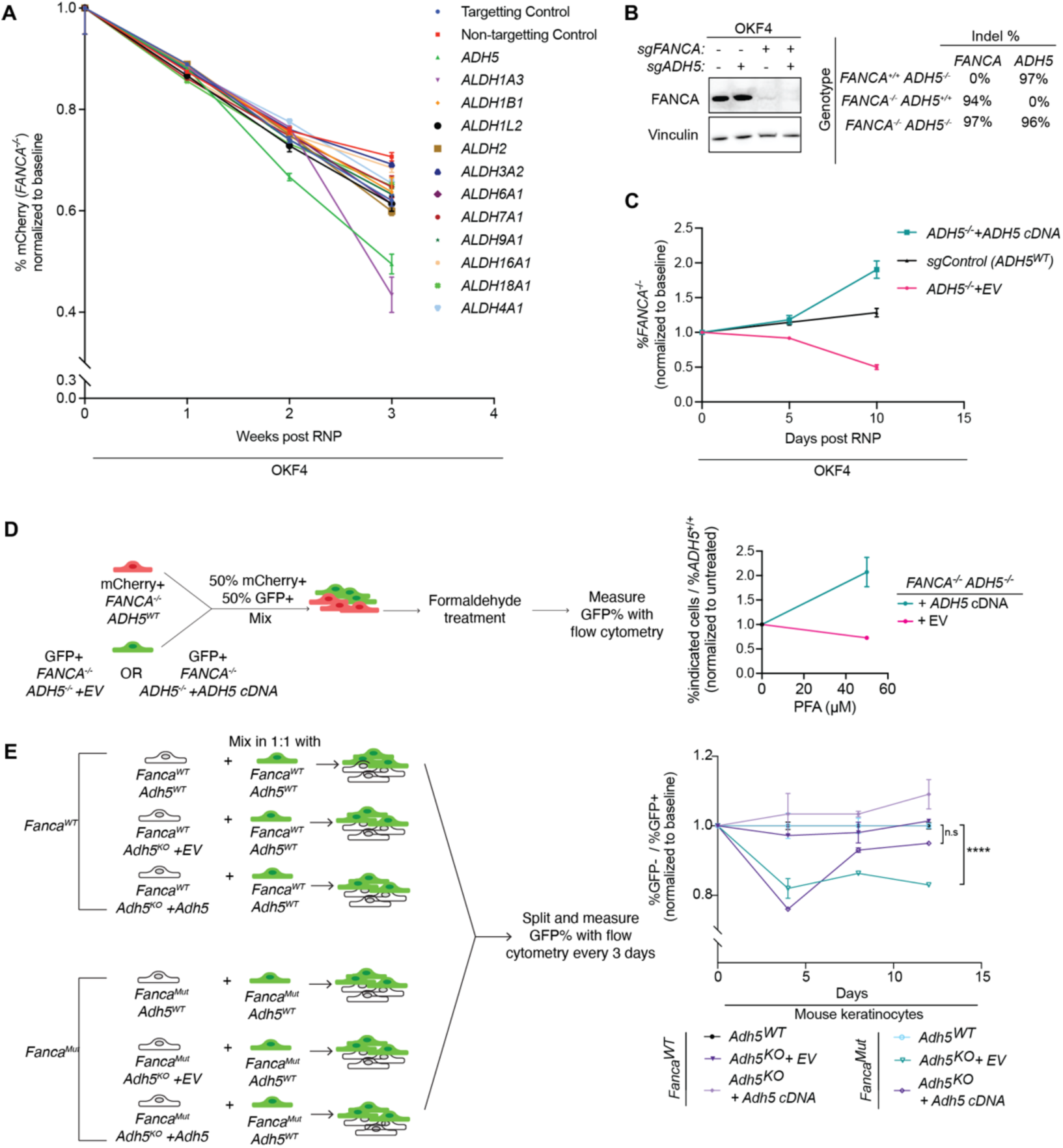
The loss of ADH5 leads to a growth defect in FA-deficient keratinocytes and sensitizes them to formaldehyde. (A) Time course of the percent of FA-deficient cells over the duration of the competition assay shown in Figure 2B for the single-gene knockouts of *ADH* or *ALDH* in OKF4. (B) Validation of *FANCA* and *ADH5* knockout in OKF4 cells. *Left*: Immunoblotting of FANCA in OKF4 cells with CRISPR-Cas9 knockouts with indicated sgRNAs. Vinculin was used as a loading control. *Right*: Summary of the percent of insertion and deletion (indel) at each gene locus. (C) Time course of the percentage of FA-deficient cells corresponding to the competition assay shown in Figure 2D. (D) Sensitivity to PFA in OKF4 *ADH5* knockout cells complemented with either empty vector (EV) or *ADH5* cDNA, measured by competition assay. *Left*: Schematic of competition assay setup. Cells were treated with indicated dose of PFA and cultured for 3 days before they were analyzed by flow cytometry. *Right*: The ratio of *FANCA^−/−^ ADH5^−/−^* over *FANCA^−/−^* at each dose was normalized to the untreated control. Data are presented as mean ± SD. (E) Competition assay in mouse keratinocytes with *Adh5* knockout as indicated in the schematic (left). Time course of the percentage of GFP-cells over the duration of the experiment corresponding to Figure 2E.

**Supplemental Figure 3.**
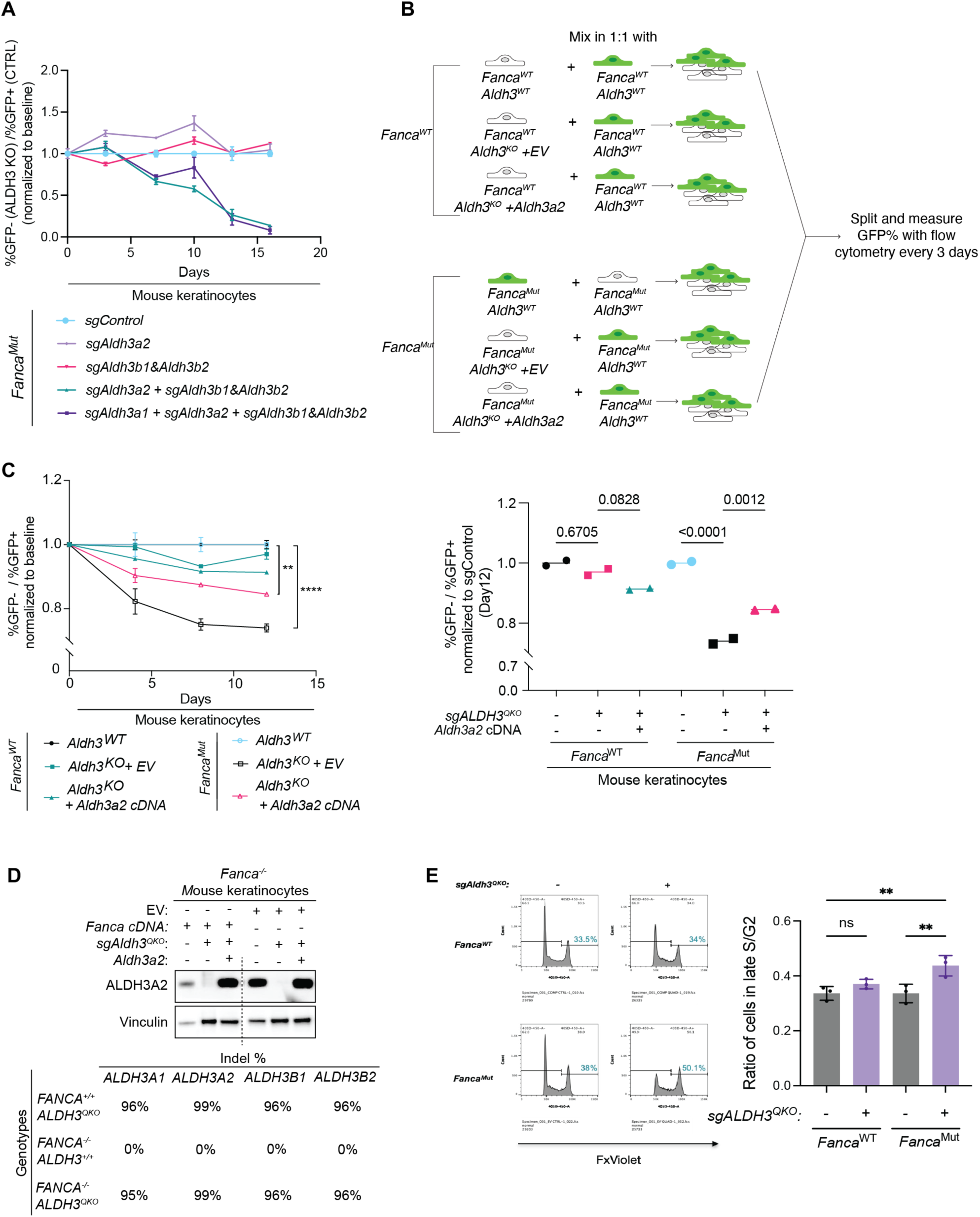
The loss of ALDH3 family genes leads to a growth defect in FA-deficient keratinocytes and sensitizes them to treatment with inducer of lipid peroxidation. (A) Time course of the percent of GFP-cells corresponding to the competition assay in Figure 3D. (B) Schematic of competition assay setup for mouse keratinocytes with *Aldh3* quadruple knockout. (C) Competition assay in mouse keratinocytes with *Aldh3* quadruple knockout in FA-competent (*Fanca^WT^*) or FA-deficient (*Fanca^Mut^*) backgrounds. For each sample, the ratio of %GFP-over %GFP+ cells after 12 days of culture was normalized to the starting ratio at the beginning of experiment and then to sgControl. *Left*: Time course of the percent of GFP-cells corresponding to the competition assay shown in Supp Figure 3B. Each dot represents one technical replicate, and data are presented as mean ± SD. *Right*: Quantification of competition assay after 12 days of culture. Each dot represents one technical replicate, and statistical significance was evaluated by one-way Kruskal-Wallis ANOVA (D) Validation of *Aldh3* quadruple knockout in mouse keratinocytes. *Top*: Immunoblotting of ALDH3A2 in mouse keratinocytes with CRISPR-Cas9 knockouts with the indicated sgRNAs. Vinculin was used as a loading control. *Bottom*: Summary of the percentage of insertion and deletion (indel) at each gene locus. (E) Cell cycle analysis in *Aldh3* quadruple knockouts in FA-competent and FA-deficient mouse keratinocytes. *Left*: Representative FxViolet DNA content histograms. *Right*: Quantification of percentage of cells in late S/G2 phase. Each dot represents one technical replicate, and statistical significance was evaluated by one-way Kruskal-Wallis ANOVA.

**Supplemental Figure 4.**
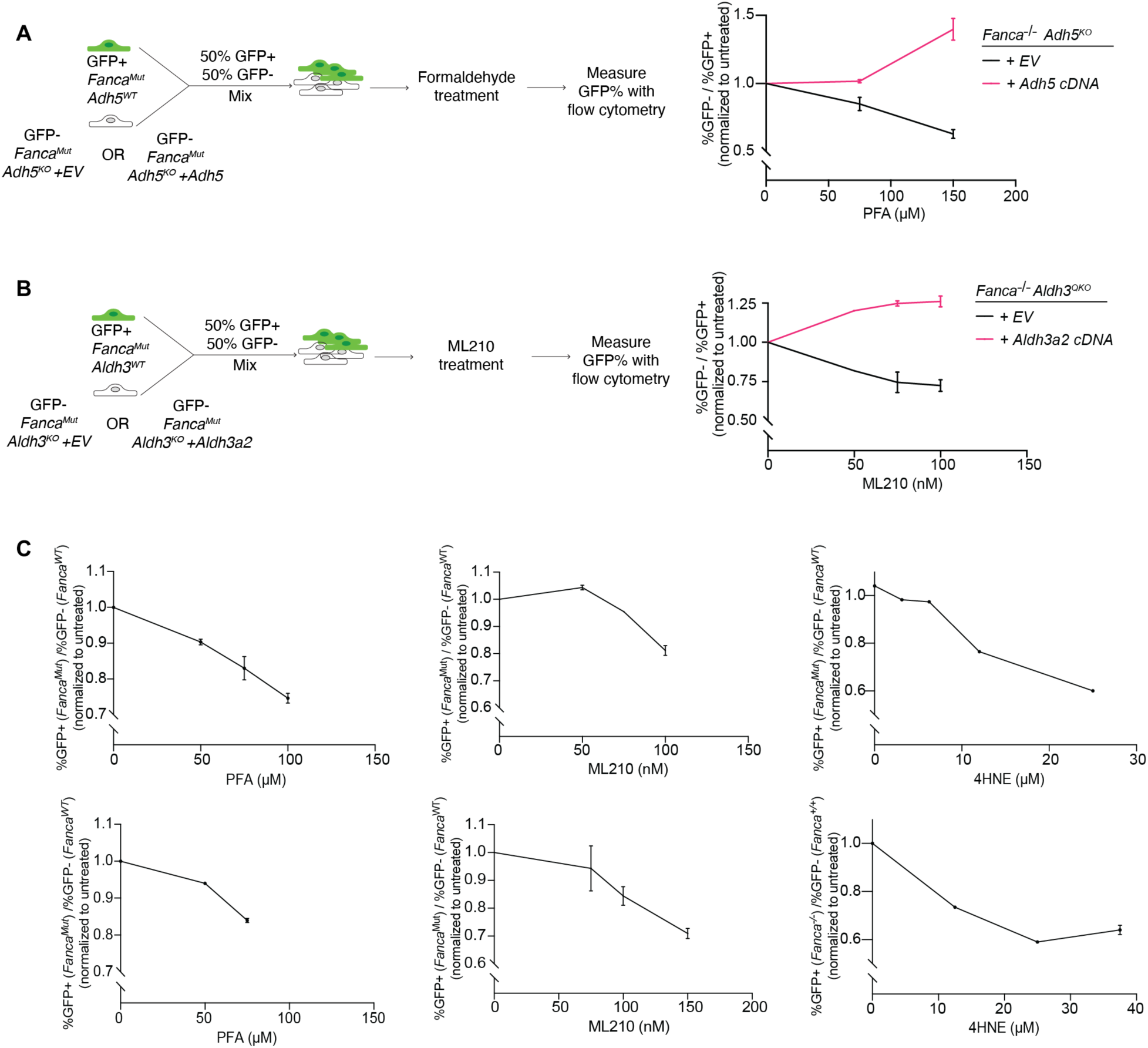
FA-deficient mouse keratinocytes exhibit sensitivity to multiple inducers of ICLs and ferroptosis. (A) Sensitivity to PFA in mouse keratinocyte *Adh5* knockout complemented with either empty vector (EV) or *Adh5* cDNA, measured by competition assay. *Left*: Schematic of competition assay setup. Cells were cultured with indicated dose of PFA and cultured for 3 days before they were analyzed by flow cytometry. *Right*: Ratio of percentage of GFP-over percentage of GFP+ cells were normalized to the untreated control. Data are presented as mean ± SD. (B) Sensitivity to ML210 in mouse keratinocyte *Aldh3* quadruple knockout complemented with either empty vector (EV) or *Aldh3a2* cDNA, measured by competition assay. *Left*: Schematic of competition assay setup. Cells were treated with indicated dose of ML210 and cultured for 3 days before they were analyzed by flow cytometry. *Right*: Ratio of percentage of GFP-over percentage of GFP+ cells were normalized to the untreated Data are presented as mean ± SD. (C) Replicate competition assays assessing the sensitivity of FA-deficient mouse keratinocytes treated with increasing doses of PFA, ML210, or 4HNE, compared to FA-competent controls. Cells were treated and cultured for 3 days prior to flow cytometry analysis. *Fanca^Mut^* cells exhibited increased sensitivity to PFA, ML210, and 4HNE.

**Supplemental Figure 5.**
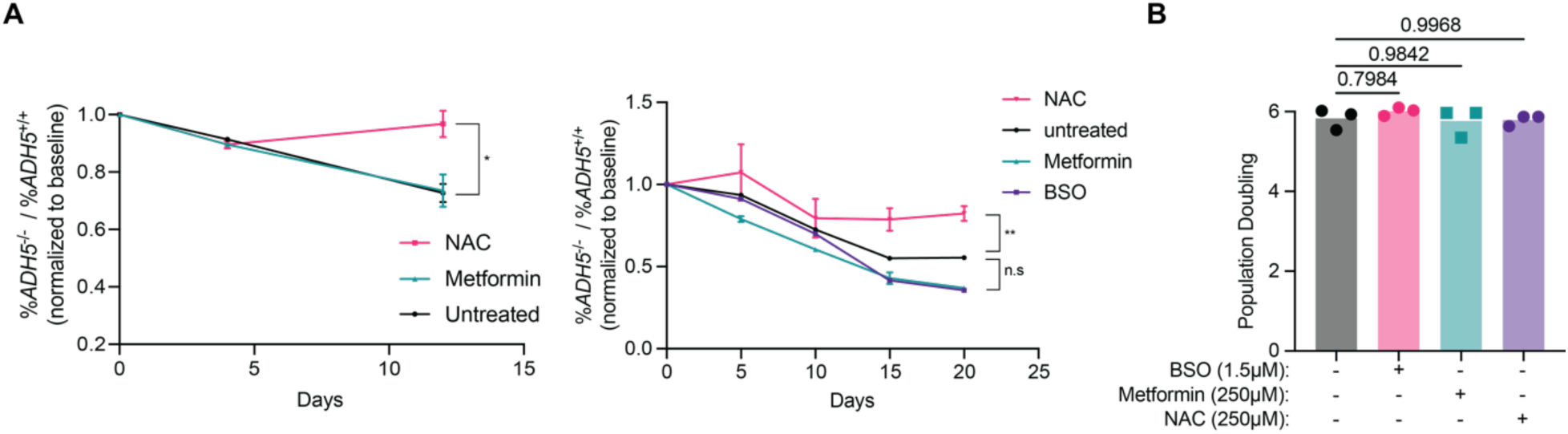
The growth defect of FA-deficient *ADH5^−/−^* OKF4 is partially rescued with supplementation of NAC, but not metformin. (A) Competition assays assessing the growth of FA-deficient *ADH5^−/−^* compared to FA-deficient *ADH5^+/+^* OKF4 cells, treated with the indicated doses of NAC, metformin or BSO. Cells were cultured for the indicated number of days, and treated once daily with 250µM NAC, 250µM metformin, and 1.5µM BSO. Only NAC treatment partially rescued the dropout of FA-deficient *ADH5^−/−^* OKF4 cells. (B) Growth of *FANCA^−/−^* cells measured by cumulative population doublings over 8 days with daily treatment of NAC, metformin, and BSO.

**Table S1.**
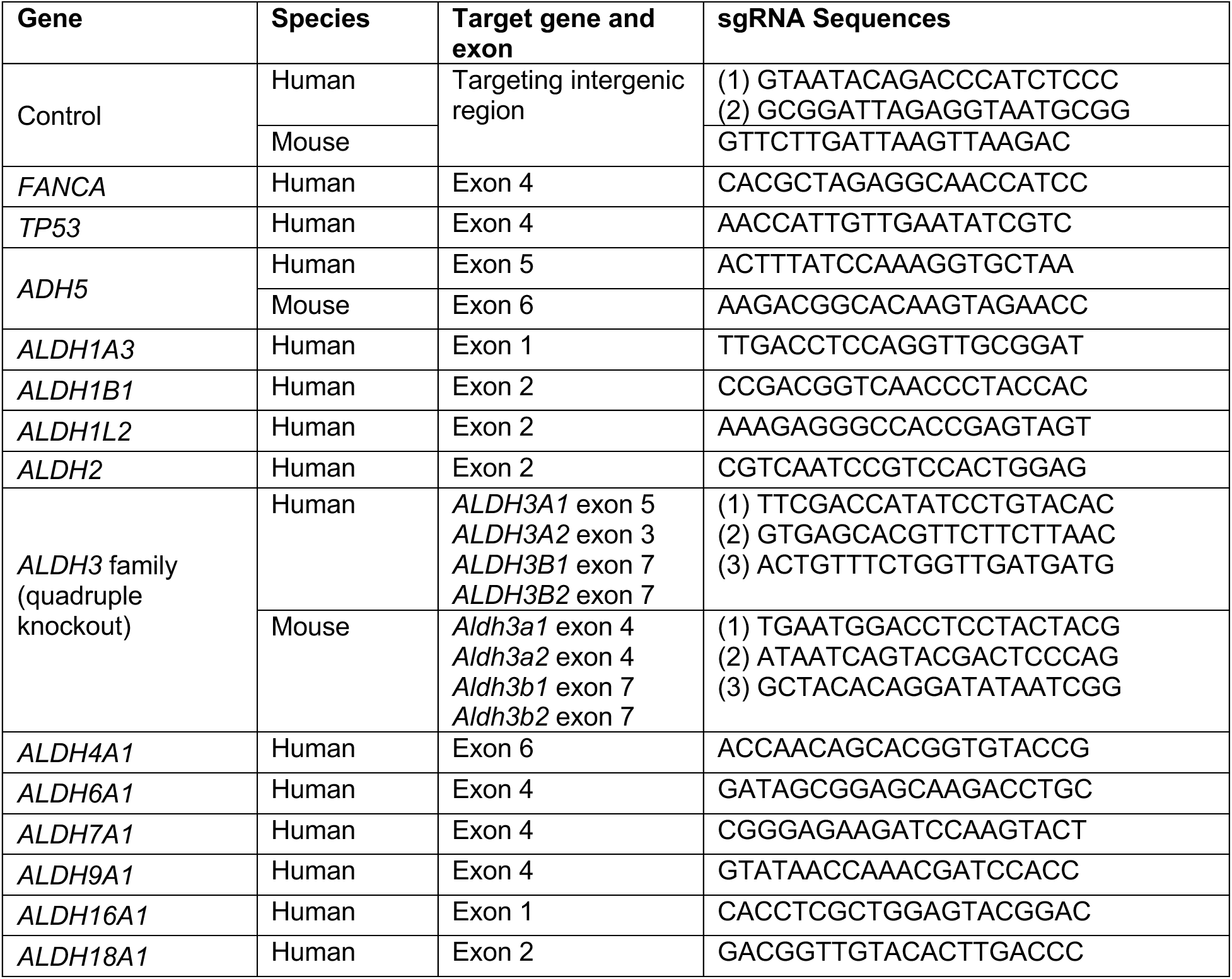
Sequences of sgRNAs.

**Table S2.**
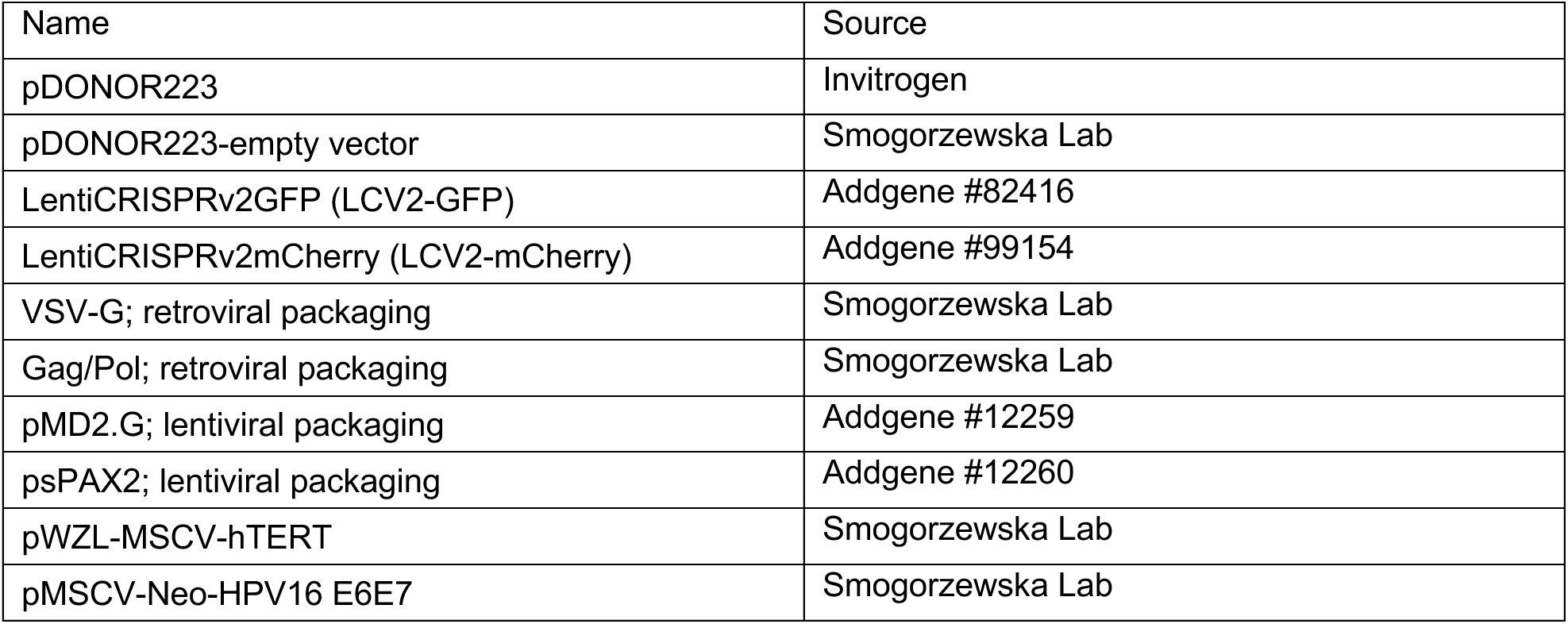
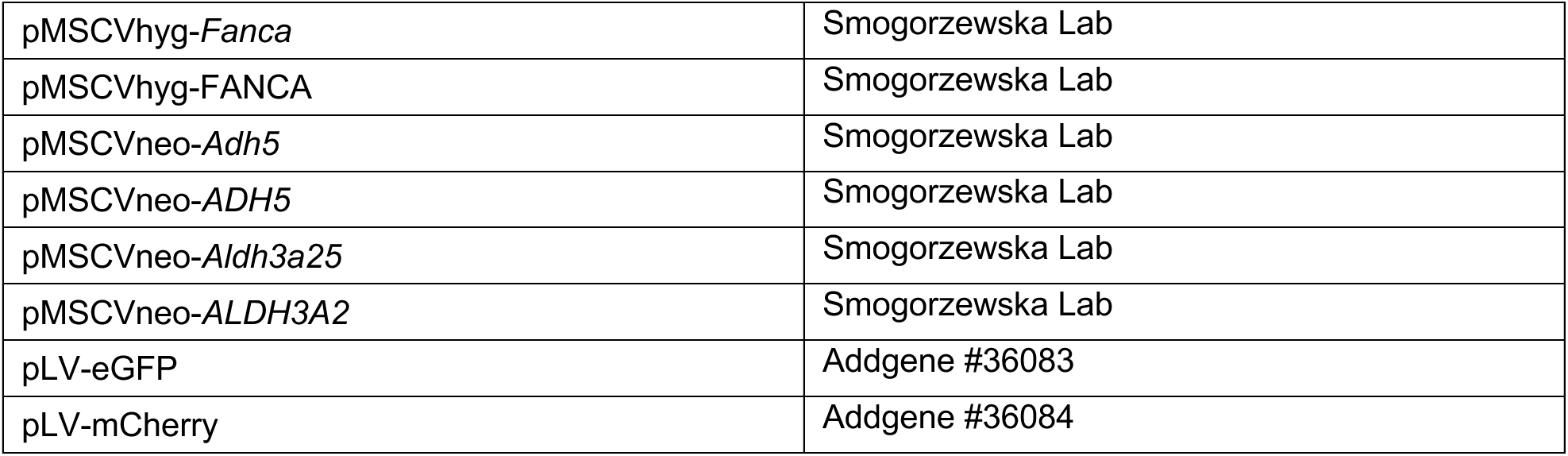
Plasmids used in this study.

